# Characterization of acetate catabolism in *Chlamydomonas reinhardtii* reveals distinct roles for *ACS1* and *ACK2* in regulating cell growth and carbon storage

**DOI:** 10.64898/2026.06.25.734523

**Authors:** Abdulmajid M. Alrefaie, Yi-Ying Lee, Yantao Li

## Abstract

Acetate metabolism drives mixotrophic and heterotrophic growth in some microalgae. Acetyl-CoA synthetase (ACS) and acetate kinase (ACK) are often considered the main enzymes involved in acetate catabolism in microalgae; however, their contributions to metabolic flux and carbon allocation are not fully understood. In this study, the functions of cytosolic *ACS1* and mitochondrial *ACK2* were characterized using two knockout mutants of the model microalga *Chlamydomonas reinhardtii*. The *acs1* mutant exhibited a growth-oriented phenotype, characterized by 29.8% faster cell growth at 96 h and up to a 15.5% higher acetate depletion rate, yet showed a 38.3% lower triacylglycerol (TAG) content at 48 h under heterotrophic conditions. By contrast, the *ack2* mutant exhibited an altered carbon-allocation phenotype under heterotrophic conditions. Despite an up to 32.4% lower respiratory oxygen consumption rate and a 27.7% reduction in cell density, *ack2* exhibited a 39.3% higher biomass concentration and a 90.4% greater dry weight per cell than the wild type at 96 h. Biochemical analysis revealed that *ack2* accumulated 23.3% more carbohydrate than the wild type at 120 h under heterotrophic conditions, whereas its TAG level remained comparable to that of the wild type. These findings suggest that, under heterotrophic conditions, the loss of cytosolic *ACS1* facilitates cell growth and division at the expense of TAG biosynthesis, whereas the loss of mitochondrial *ACK2* regulates growth by affecting carbon flux toward biomass and carbohydrate accumulation. This work provides insight into acetate catabolism in *C. reinhardtii* and suggests targets for engineering microalgae for production of biomass and bioproducts.

## 1. INTRODUCTION

Microalgae are photosynthetic microorganisms, key contributors to the global carbon cycle, and have been used as feedstocks for production of biofuels and bioproducts [1, 2]. Their ability to efficiently harness energy from light or organic compounds, coupled with remarkable metabolic plasticity, allows them to adapt to varying environments through autotrophic, mixotrophic, or heterotrophic growth modes [3, 4]. Heterotrophy enables growth in the dark using organic carbon substrates, a versatility that positions microalgae as robust platforms for various biotechnological applications [5, 6].

Historically, industrial microbial cultivation has relied on heterotrophic fermentation metabolizing simple organic carbon sources, such as glucose and acetate [7, 8]. This approach has been adapted for microalgae, such as *Crypthecodinium cohnii*, for commercial omega-3 fatty acids production [6, 9]. Heterotrophic cultivation offers advantages for industrial scalability, including fast growth rate, ultrahigh cell density, non-self-shading, lower contamination risk, and standardized control over environmental parameters [6, 8, 9].

Among model organisms, the green microalga *Chlamydomonas reinhardtii* is able to utilize acetate as an organic carbon source [10]. Unlike many industrial heterotrophs that utilize glucose, *C. reinhardtii* lacks hexose transporters and relies on acetate to fuel central carbon metabolism that drives growth and essential metabolic processes [11, 12]. Acetate supplementation has been shown to suppress photosynthesis [12], redirecting carbon flux toward lipid storage [13]. The extensive genomic [11, 14] and genetic resources [15–17] available for *C. reinhardtii* make it a powerful model system to study acetate metabolism and lipid biosynthesis.

Acetate catabolism generates acetyl-CoA, a central precursor for both energy production and lipid biosynthesis [17, 18]. In *C. reinhardtii*, this conversion is mediated primarily through two routes: via acetyl-CoA synthetases (ACSs) or via a two-step enzymatic reaction catalyzed by acetate kinase (ACK) and phosphate acetyltransferase (PAT) [18, 19]. These enzyme families include multiple isoforms, some of which differ in subcellular localization and metabolic function. For example, ACS1 is cytosolic, supporting cytosolic acetyl-CoA pools, whereas ACK2 is mitochondrial, feeding mitochondrial acetyl-CoA into the tricarboxylic acid (TCA) cycle [17, 20]. This spatial organization likely allows *Chlamydomonas* cells to tune carbon flux between the energy-generating TCA cycle and fatty acid biosynthesis [19]. However, the contributions of each pathway to acetate catabolism and their impacts on downstream pathways remain elusive.

In this study, we characterized *acs1* and *ack2* mutants to assess how disruption of *ACS1* and *ACK2*, respectively, affects growth, respiration, and carbon allocation under mixotrophic and heterotrophic conditions. By combining measurements of growth, nutrient consumption, photosynthetic efficiency, respiration, and storage compound accumulation, we investigated how *Chlamydomonas* cells utilize acetate-derived carbon for growth and storage. Insights into these metabolic pathways improve our understanding of the mechanisms underlying acetate catabolism in algae and provide a foundation for engineering strains capable of converting low-cost organic carbon feedstocks into renewable fuels and high-value bioproducts.

## 2. MATERIALS AND METHODS

### 2.1. Strain and culture conditions

The *C. reinhardtii* strains used in this study were the cell wall-deficient wild type (CC-5325), the acetyl-CoA synthetase mutant *acs1* (LMJ.RY0402.072886), and the acetate kinase mutant *ack2* (LMJ.RY0402.122457). All strains were obtained from the CLiP library (Chlamydomonas Resource Center) [21] and maintained on Tris–acetate–phosphate (TAP) agar plates. The *acs1* and *ack2* mutant strains were maintained in TAP medium supplemented with paromomycin. For experimental inoculum cultures, single colonies were inoculated into 10 mL of liquid TAP and incubated for 5–7 days at 25°C with continuous shaking under constant illumination to reach the exponential phase. Experimental cultures were initiated in 100 mL of fresh TAP medium, with inoculation of 10 mL of seed culture at a starting cell density of ca. 5.8 × 10⁵ cells mL⁻¹.

Cultures were grown under two conditions: mixotrophic cultures were maintained under continuous illumination at 50 μmol photons m⁻² s⁻¹ at 25°C, whereas heterotrophic cultures were maintained in complete darkness (0 μmol photons m⁻² s⁻¹) at 25°C. All cultures were grown with continuous shaking at 150 rpm. TAP medium (pH 7.0) was prepared according to the recipe of Gorman and Levine (1965). For solid TAP medium, 15 g L⁻¹ agar was added. All cultures were routinely checked for bacterial contamination using Reasoner’s 2A Agar Broth (R2B). All experiments were set up in biological triplicates.

### 2.2. Growth and biomass measurements

Growth parameters were measured every 24 h for 6 days. Cell density was determined by counting with a hemocytometer. Optical density was measured at 750 nm (OD₇₅₀) with a NanoDrop 2000c spectrophotometer (Thermo Fisher Scientific, Danvers, MA) to minimize interference from chlorophyll pigments [22, 23]. Cell size distributions were analyzed with a Multisizer 4e Coulter Counter (Beckman Coulter, Inc, Indiana, USA).

Biomass was determined gravimetrically. At each time point, a 10 mL culture was filtered under vacuum through a pre-weighed VWR 696 glass microfiber filter, then dried in an oven at 100°C for 24 h and weighed on an analytical balance (Sartorius AG, Göttingen, Germany).

### 2.3. Nutrient consumption analysis

The concentrations of acetate, ammonia, and phosphate in heterotrophic cultures were measured. At each time point, 1.5 mL of culture was collected and centrifuged at 13,000 × g for 10 min at 25 °C. The collected supernatant was filtered through a 0.22 μm nylon membrane prior to analysis.

Acetate concentration was analyzed by high-performance liquid chromatography (HPLC) using an Agilent 1260 Infinity II system (Agilent Technologies, Santa Clara, CA). Separation was performed using a Sepax Carbomix H-NP5 column, 4.6 × 300 mm (Sepax Technologies, Inc., Newark, DE) with 25 mM sulfuric acid as the mobile phase at a flow rate of 0.25 mL min⁻¹. A standard curve was constructed using serial dilutions of sodium acetate as the reference standard to quantify acetate in experimental samples.

Ammonia was quantified using a HACH DR300 Colorimeter (Hach Company Inc., Loveland, CO) and the high-range Salicylate Test ‘N Tube method, following the manufacturer’s instructions [24]. Absorbance was measured at 655 nm, and ammonia concentrations were determined by comparing sample values against a standard curve generated from ammonium chloride solutions of known concentrations.

Phosphate concentrations were measured using a malachite green assay. The assay involved combining the sample (with hydrolyzed phosphate) with assay reagents, incubating for 20 min, and measuring absorbance at 595 nm against a 0–500 μM KH₂PO₄ standard curve with a SpectraMax M5 microplate reader (Molecular Devices, San Jose, CA) [25].

### 2.4. Mutant confirmation

Genomic DNA was extracted using the E.Z.N.A. Plant & Fungal DNA Kit (Omega Bio-tek, Norcross, GA). Gene sequences were retrieved from Phytozome (v5.6) and NCBI (v5.5).

Mutations in the *ACS1* and *ACK2* genes were confirmed by PCR. Flanking primer pairs that match genomic regions surrounding the predicted insertion sites and insertional primers (oMJ913 or oMJ944) that match the insertion cassette (oMJ913 or oMJ944) were used, consisting of one genomic flanking primer and one primer specific to the inserted DNA cassette [21]. PCR amplifications were performed using Phusion High-Fidelity DNA Polymerase (Thermo Scientific, Waltham, MA) and Taq DNA Polymerase (New England Biolab, Ipswich, MA).

PCR products were sequenced to confirm the exact insertion sites within *ACS1* and *ACK2*. These approaches followed established methods for verifying insertional mutants in *C. reinhardtii* [21].

### 2.5. Photosynthetic efficiency and respiration measurements

The maximum quantum efficiency of Photosystem II (PSII), known as *Fv/Fm*, was measured using a WALZ PAM-2500 fluorometer (Heinz Walz GmbH, Effeltrich, Germany). Samples were dark-adapted for 20 min before measurement to allow oxidation of PSII reaction centers. *Fv/Fm* was calculated as (Fₘ − F₀)/Fₘ. Cellular respiration rates were measured in real time using a Resipher 96-well plate reader system (Lucid Scientific, Atlanta, GA). Strains were inoculated into 96-well plates containing 200 μL TAP medium. The instrument measured oxygen gradients generated by cells and reported the average respiratory oxygen consumption rate (OCR, fmol mm⁻² s⁻¹) and dissolved oxygen concentration (μM).

### 2.6. Protein localization prediction

Subcellular localization predictions were generated using PredAlgo [26], and DeepLoc 2.0 [27]. The predicted localizations were compared with experimental data reported in the literature [17].

### 2.7. Total lipid and total TAG quantification

Total lipids were extracted from approximately 100 mg of freeze-dried biomass using a FreeZone Benchtop Freeze Dryer (Labconco Corporation, Kansas City, MO), followed by extraction with chloroform:methanol (2:1, v/v). Phase separation was induced by adding water to a final ratio of chloroform:methanol:water (2:1:0.75, v/v/v), followed by centrifugation. The organic phase was collected, and the extraction was repeated twice to maximize lipid recovery. Extracts were dried under nitrogen and quantified gravimetrically. Lipid contents were normalized to biomass dry weight and cell density.

For TAG separation, total lipid extracts were separated by thin-layer chromatography (TLC) on silica gel 60 F₂₅₄ aluminum-backed plates (Supelco-MilliporeSigma, Darmstadt, Germany) using a solvent system of hexane: t-butyl methyl ether: acetic acid (80:20:2, v/v/v). Plates were visualized with a copper(II) sulfate pentahydrate spray, followed by charring. TAG bands were identified by comparison with those from TAG standards.

For quantification, TAG bands separated by TLC were scraped from the silica plate and subjected to transesterification with 1% sulfuric acid in methanol at 85°C for 1.5 h in the presence of an internal standard (C17:0). The resulting fatty acid methyl esters (FAMEs) were extracted with hexane and analyzed by gas chromatography–mass spectrometry (GC–MS) using a Trace 1310 gas chromatograph coupled to a TSQ 8000 triple quadrupole mass spectrometer (Thermo Fisher Scientific, Waltham, MA, USA).

### 2.8. Total carbohydrate quantification

Total carbohydrate content was measured from 10 mg of freeze-dried biomass using a phenol-sulfuric acid colorimetric assay with glucose as the calibration standard. The biomass was first treated with concentrated acetic acid and acetone to remove pigments and then hydrolyzed by boiling in 4 M trifluoroacetic acid for 4 h. After dilution and clarification by centrifugation, aliquots were reacted with freshly prepared phenol-sulfuric acid reagent and heated for 20 min to develop the colorimetric reaction. Absorbance was measured at 490 nm, and carbohydrate concentration was determined from the standard curve made with glucose standards of known concentrations.

### 2.9. Statistical analysis

All data points were presented as the mean of three biological replicates (n = 3), with error bars representing the standard deviations (SD). Statistical significances between wild-type and mutant strains were evaluated using a two-tailed Student’s *t*-test, with a statistical significance indicated by * (*t*-test, *p* < 0.05).

## 3. RESULTS AND DISCUSSION

### 3.1. Molecular confirmation of acs1 and ack2 insertion mutants

As the putative *acs1* and *ack2* mutants were obtained from the CLiP library [21], we first confirmed the mutations in the *acs1* and *ack2* strains at the genomic DNA level using antibiotic resistance (Fig. S1), PCR genotyping (Fig. S2), and Sanger sequencing, which confirmed insertions in exon 17 of *ACS1* and intron 1 of *ACK2* (Fig. S3). RT-qPCR analysis verified the transcript-level disruption in *acs1*, showing undetectable *ACS1* expression (Fig. S4). For confirmation of the *ack2* strain, a near full-length *ACK2* cDNA fragment (1919 bp) was detected in the wild type CC-5325 by RT-PCR amplification, but was not detectable in the *ack2* strain.The absence of a full-length *ACK2* transcript infers that the *ACK2* gene is disrupted (Fig. S5).

### 3.2. Growth and biomass accumulation of acs1 and ack2

*C. reinhardtii* possesses a spatially organized acetate metabolism with compartment-specific isoforms: ACS1 is predicted to be cytosolic, supporting cytosolic acetyl-CoA pools, whereas its counterparts ACS2 and ACS3 are predicted to be localized to the plastid and mitochondria, respectively (Supplementary Table 1). Previous experimental work has confirmed that ACK1 is localized to the plastid and ACK2 is mitochondrial (Supplementary Table 1), feeding acetate-derived acetyl-CoA into the TCA cycle [17, 20].

Next, the *acs1* and the *ack2* strains were grown under mixotrophic and heterotrophic conditions. Under mixotrophic conditions, the *acs1* strain showed a clear growth advantage. At 96 h, the *acs1* strain reached 8.1×10⁶ cells mL⁻¹, which was 65.2% higher (*t*-test, *p* < 0.05) than the wild-type strain (CC-5325) of 4.9×10⁶ cells mL⁻¹, and achieved an OD₇₅₀ of 2.5, which was 25.1% higher than the control (Fig. 1a; 1b). On the contrary, the *ack2* strain grew significantly more slowly. At 96 h, the cell density of the *ack2* strain reached 4.2×10⁶ cells mL⁻¹, which was about 14.1% lower than the wild-type strain (*t*-test, *p* < 0.05), and OD₇₅₀ was 1.9, which was 5.1% lower than the wild-type (Fig. 1a; 1b).

**Figure 1.**
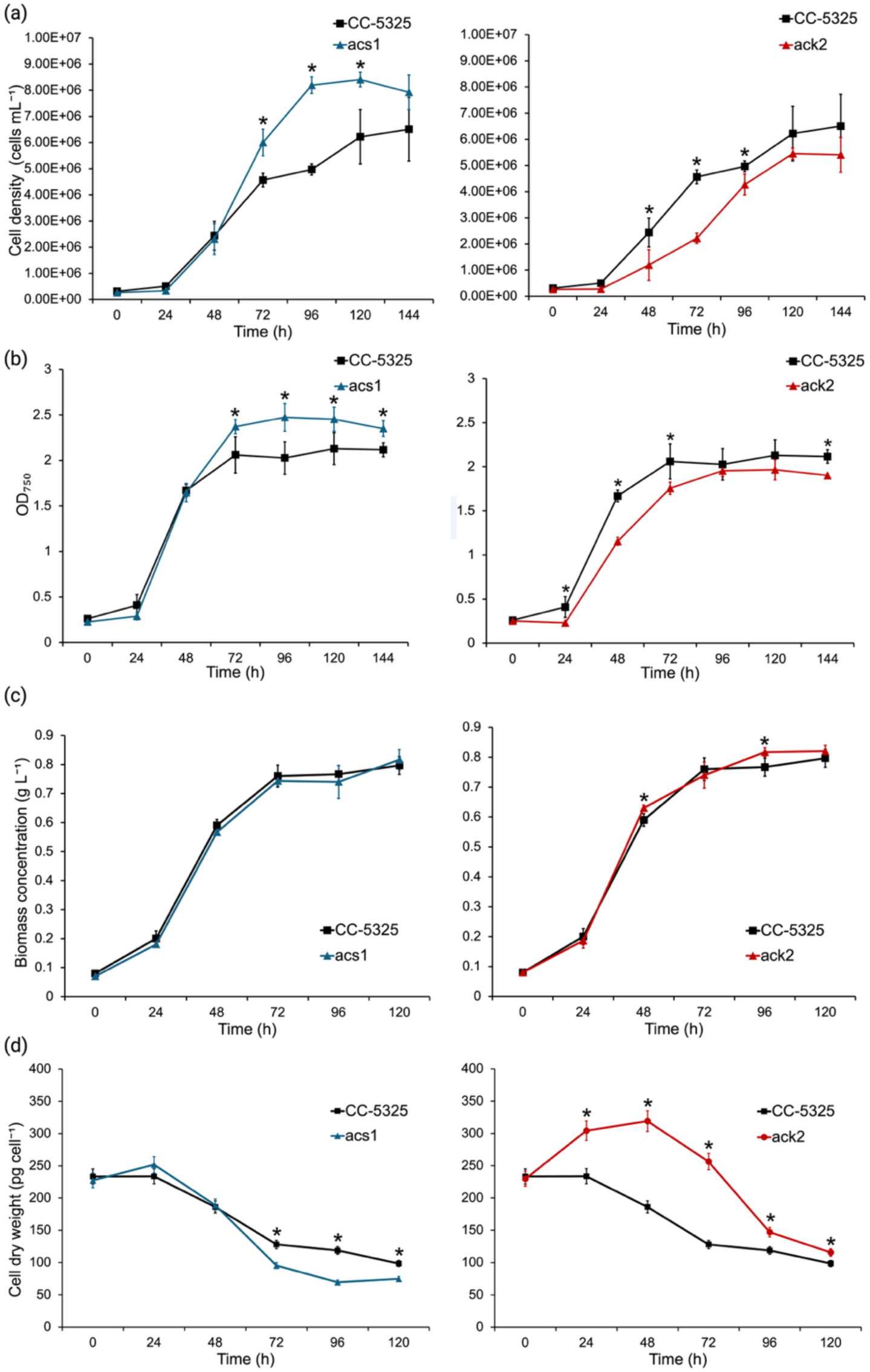
Growth dynamics, biomass accumulation, and cell dry weight of the *acs1* and *ack2* mutants under mixotrophic conditions. From top to bottom, each panel showed cell density (a, cells mL⁻¹), optical density (b, OD_750_), biomass concentration (c, g L⁻¹), and dry weight per cell (d, pg cell⁻¹) for *acs1* (blue triangle), *ack2* (red circle), and CC-5325 (black square). The left panels showed measurements for *acs1* compared with CC-5325, and the right panels showed measurements for *ack2* compared with CC-5325. Biological triplicate cultures were maintained in TAP medium at 25 °C under continuous light at 50 μmol photons m^−2^ s^−1^ (mixotrophic growth). Error bars represented standard deviations (n=3). Statistical significances between CC-5325 and the mutant strains were determined using a two-tailed Student’s *t*-test, with asterisks (*) indicating a *p*-value < 0.05.

Under heterotrophic conditions, similar phenomena were found. The *acs1* mutant maintained its growth advantage, peaking at 6.1×10⁶ cells mL⁻¹ at 96 h and was 29.8% higher (*t*-test, *p* < 0.05) than the wild-type at 4.7 × 10⁶ cells mL⁻¹ (Fig. 2a). Optical density for the heterotrophic *acs1* culture was also consistently higher than the control across the stationary phase (Fig. 2b). In contrast, the *ack2* mutant showed an apparent growth defect in the dark. At 96 h, the *ack2* mutant reached only 3.4 × 10⁶ cells mL⁻¹, which was 27.7% lower than the wild-type (*t*-test, *p* < 0.05) with an OD₇₅₀ of 1.1, compared to 1.3 of the wild-type (Fig. 2a; 2b). These growth patterns suggest distinct roles for the *ACS1* and *ACK2* genes. The increased cell density in the *acs1* mutant suggests that loss of *ACS1* may shift acetate utilization toward cell growth. In contrast, the *ack2* mutant showed a persistently lower cell density, particularly under heterotrophic conditions where acetate serves as the sole carbon source, suggesting the importance of the *ACK2* gene to efficient acetate utilization and respiration, in line with previous findings [17, 19].

**Figure 2.**
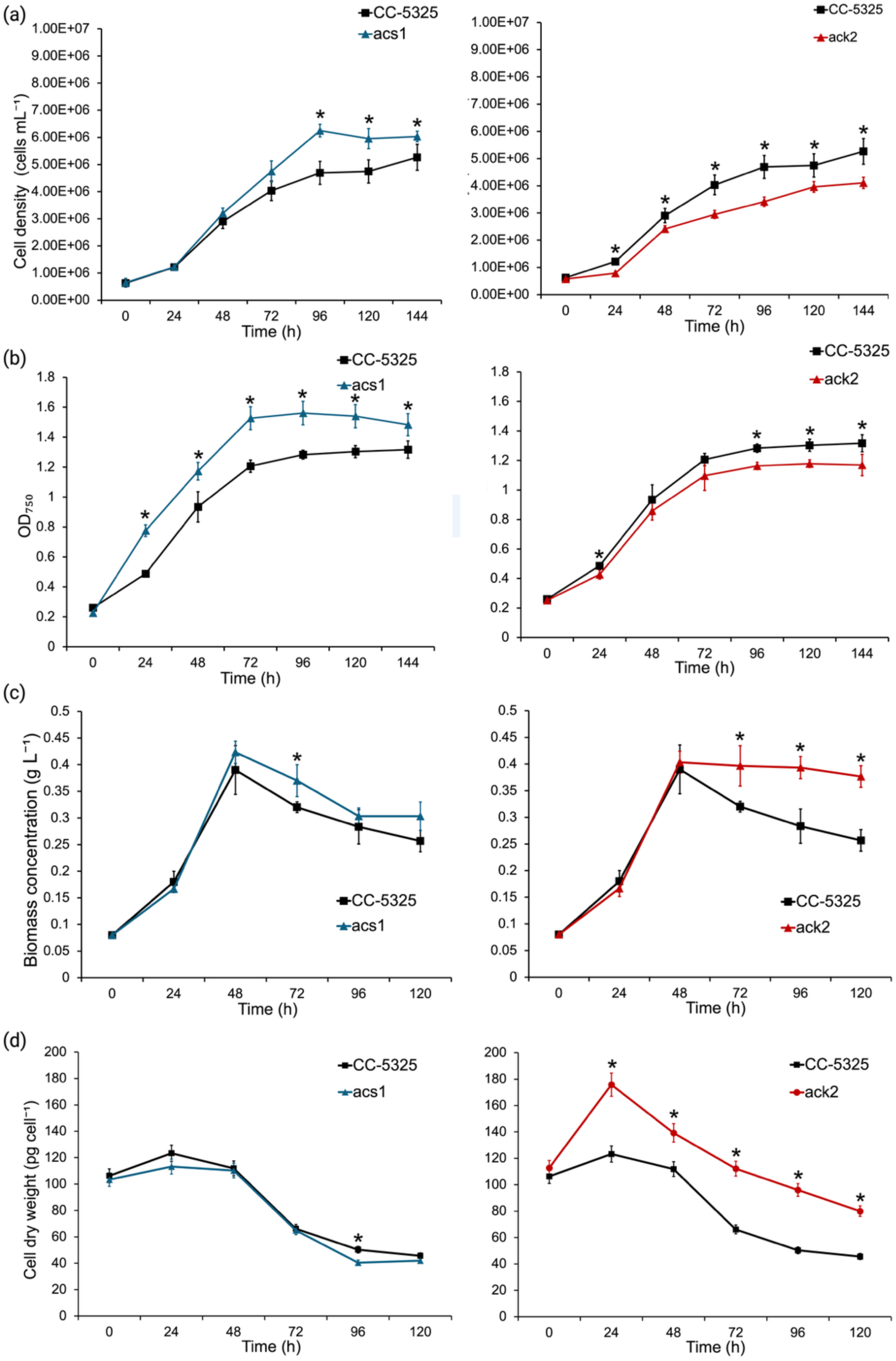
Growth dynamics, biomass accumulation, and cell dry weight of the *acs1* and *ack2* mutants under heterotrophic conditions. From top to bottom, each panel showed cell density (a, cells mL⁻¹), optical density (b, OD_750_), biomass concentration (c, g L⁻¹), and dry weight per cell (d, pg cell⁻¹) for *acs1* (blue triangle), *ack2* (red circle), and CC-5325 (black square). The left panels showed measurements for *acs1* compared with CC-5325, and the right panels showed measurements for *ack2* compared with CC-5325. Biological triplicate cultures were maintained in TAP medium at 25 °C in complete darkness (heterotrophic growth). Error bars represented standard deviations (n=3). Statistical significances between CC-5325 and the mutant strains were determined using a two-tailed Student’s *t*-test, with asterisks (*) indicating a *p*-value < 0.05.

To determine whether culture biomass reflected cell density, cell size, or both, we measured both average culture biomass concentration (g L⁻¹) and average dry weight per cell (pg cell⁻¹). Under mixotrophic conditions, the biomass concentration of the *acs1* strain was comparable to that of the wild type (Fig. 1c). However, a significantly lower dry weight of the *acs1* strain on a per cell basis was observed at 72, 96, and 120 h (*t*-test, *p* < 0.05) by 24.0%, 40.7%, and 25.0%, respectively (Fig. 1d). Similarly, the biomass density of the *ack2* strain was generally similar to the wild type under mixotrophy but slightly higher at 48 and 96 h (by 5.1% and 6.6%, respectively) (*t*-test, p < 0.05) (Fig. 1c). These differences were accompanied by significantly greater dry weight of *ack2* cells on a per cell basis by 30.6%, 71.4%, 98.4%, 20.8%, and 19.4% at 24, 48, 72, 96, and 120 h, respectively (*t*-test, *p*< 0.05) (Fig. 1d).

Under heterotrophic conditions, the *acs1* strain generally maintained biomass dry weight comparable to the wild type, except that biomass dry weight was significantly higher (*t*-test, p < 0.05) by 15.6% at 72 h, whereas dry weight per cell was reduced by 20.0% at 96 h (Fig. 2c, 2d). In contrast, *ack2* cultures showed significantly greater (*t*-test, *p*< 0.05) biomass dry weight by 23.4%, 39.3%, and 47.1%, respectively at 72, 96, and 120 h (Fig. 2c). This increase was accompanied by a significantly greater (*t*-test, *p*< 0.05) weight on a per cell basis by 45.8%, 25.0%, 69.7%, 90.4%, and 77.8%, respectively at 24, 48, 72, 96, and 120 h (Fig. 2d).

### 3.3. Nutrient consumption analysis in heterotrophic growth

Next, we monitored the concentrations of acetate, ammonia, and phosphate to determine if the growth and biomass phenotypes correlated with nutrient consumption rates. To exclude the effect of light and photosynthesis under mixotrophic conditions, they were only measured in heterotrophic cultures in which acetate was the sole carbon source. Acetate consumption of the *acs1* mutant was significantly faster than that of the wild-type. At 24 h, the acetate concentration in the *acs1* cultures was lower by 15.5%. By 48 h of heterotrophic growth, acetate (*ca.* 1.2 g L^-1^) in both mutants and the wild-type cultures was fully depleted (Fig. 3a). Comparing the *ack2* and the wild-type strains, no differences in acetate consumption were observed (Fig. 3a).

**Figure 3.**
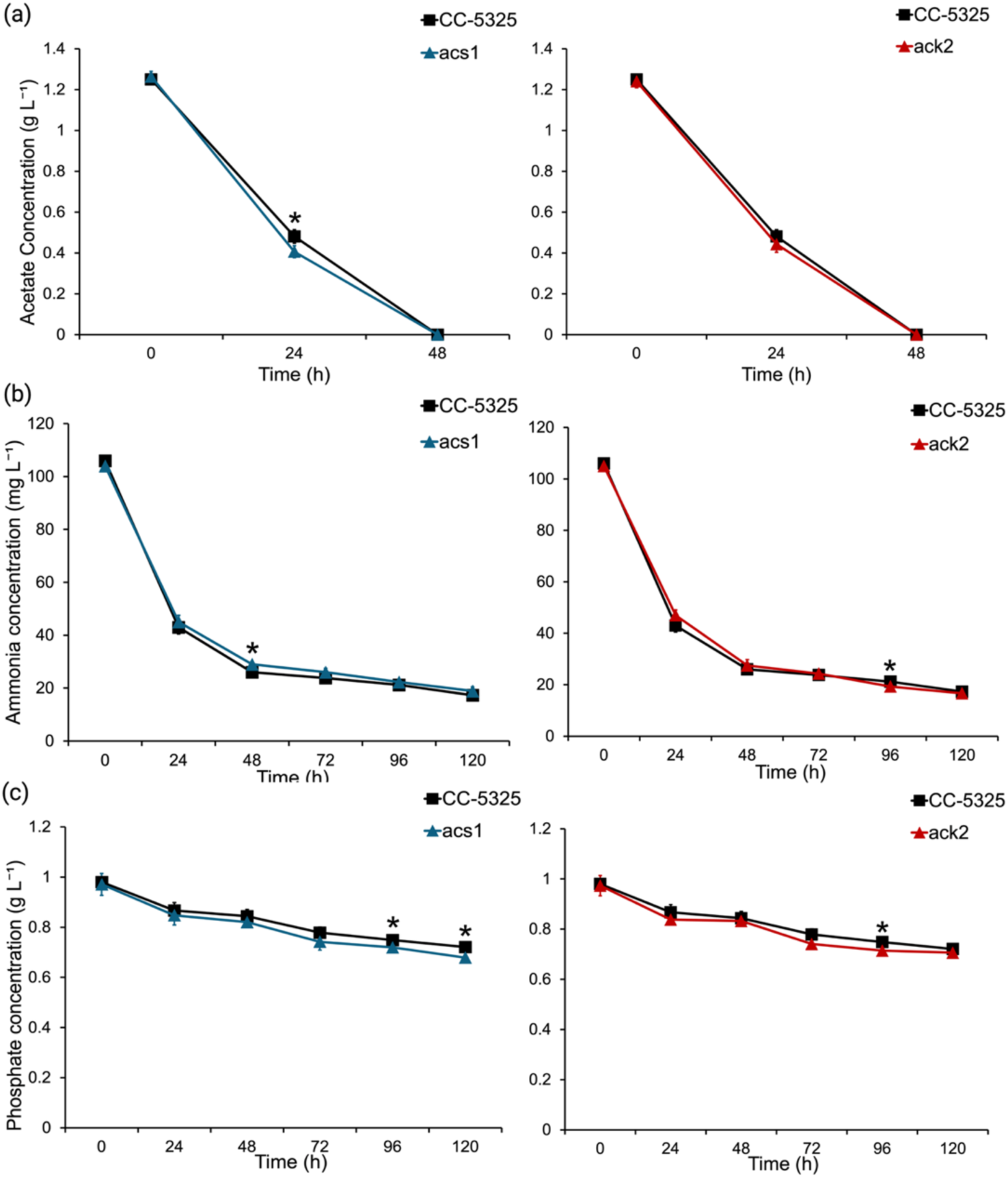
Nutrient consumption by *acs1* and *ack2* mutants under heterotrophic conditions. Concentrations of (a) acetate (g L⁻¹), (b) ammonia (mg L⁻¹), and (c) phosphate (g L⁻¹) were measured in the medium of the *acs1* (blue), *ack2* (red), and CC-5325 (black) cultures. Biological triplicate cultures were maintained in TAP medium in complete darkness at 25 °C. All concentrations were recorded every 24 h for a total of 120 h. Error bars represented standard deviations (n=3). Statistical significances between the wild-type and mutant strains were assessed using a two-tailed Student’s *t*-test, with asterisks (*) indicating a *p*-value < 0.05.

Nitrogen (as ammonia) and phosphate consumption patterns in the *acs1* and *ack2* strains were overall similar. However, the residual ammonia level in the *acs1* cultures was higher (by 11.5%) than in the wild-type cultures at 48 h, while the ammonia level in the *ack2* cultures was slightly lower (by 9.4%) than in the wild-type cultures at 96 h (Fig. 3b). For phosphate, the *acs1* and *ack2* cultures maintained slightly lower residual concentrations (by 3.9% and 4.6%, respectively) than the wild type did at 96 h (Fig. 3c).

### 3.4. Photosynthetic efficiency analysis

To assess the status of the photosynthetic apparatus, we measured maximal Photosystem II (PSII) quantum efficiency (*Fv/Fm*) in the *acs1*, *ack2*, and wild-type cultures under both mixotrophic and heterotrophic conditions. Under mixotrophic conditions, the *acs1* cultures showed signs of photosynthetic stress. *Fv/Fm* in mixotrophic *acs1* cultures declined more rapidly and was significantly lower than in the wild type from 72 h onward. It was decreased by 8.7%, 12.6%, 7.0%, and 10.3%, respectively, at 72, 96, 120, and 144 h (Fig. 4a). In contrast, *Fv/Fm* in the mixotrophic *ack2* cultures was generally similar to that of the wild-type cultures, with significant differences observed only at 48 h (*t*-test, *p* < 0.05) by an 8.9% decrease (Fig. 4a). These data suggest that disruption of *ACK2* has a limited effect on PSII efficiency under mixotrophic conditions.

**Figure 4.**
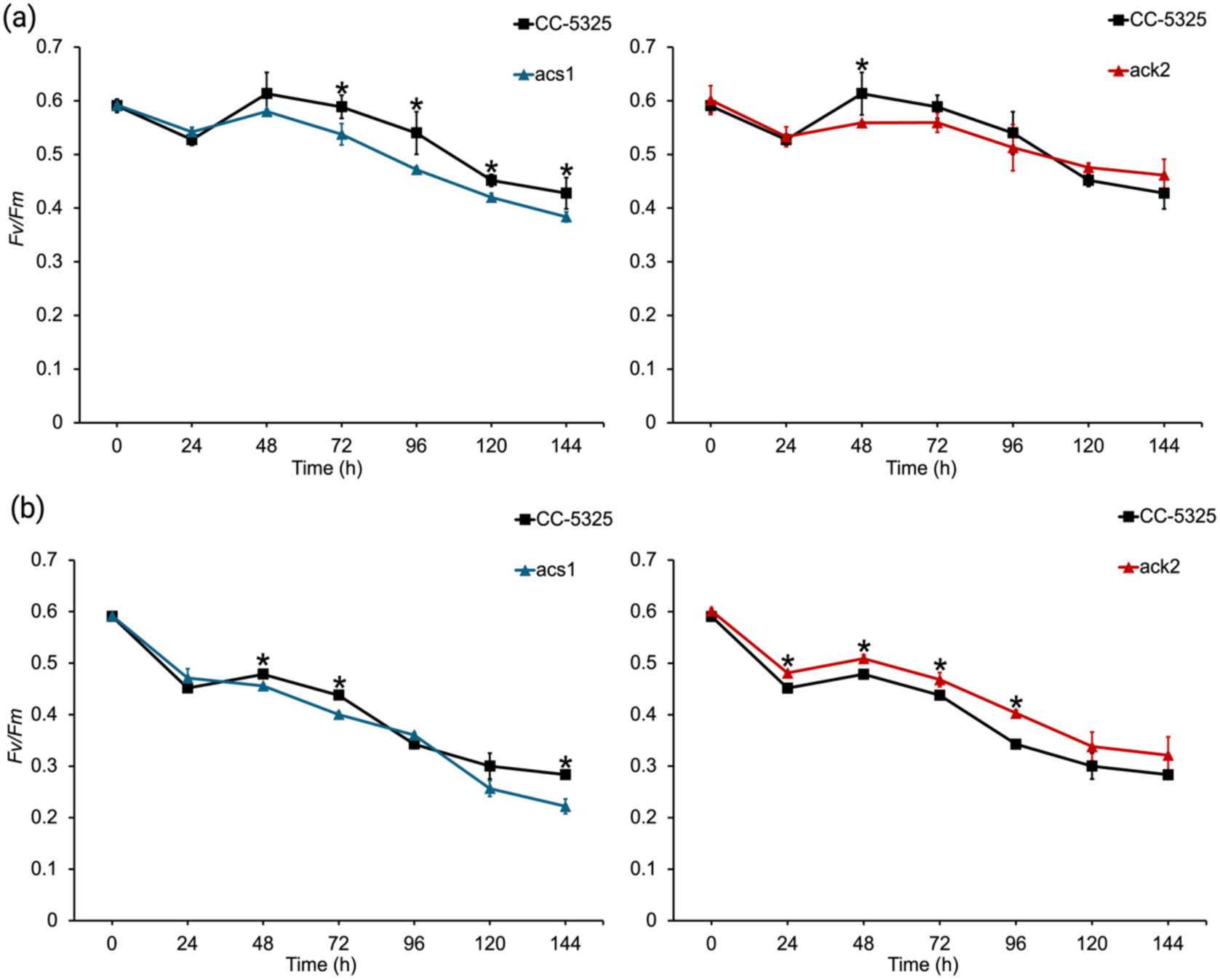
Photosynthetic efficiency (*Fv/Fm*) of the *acs1* and *ack2* mutants under mixotrophic or heterotrophic conditions. Maximum PSII quantum efficiency (*Fv/Fm*) of *acs1* (blue), *ack2* (red), and CC-5325 (black) under (a) mixotrophic and (b) heterotrophic conditions. Biological triplicate cultures were maintained in TAP medium at 25 °C. *Fv/Fm* was measured every 24 h up to 144 h. Error bars represent standard deviations (n=3). Statistical significances between CC-5325 and the mutant strains were determined using a two-tailed Student’s *t*-test, with asterisks (*) indicating a *p*-value < 0.05.

Under heterotrophic conditions, the *acs1* stress phenotype is evident. From 48 h onward, *Fv/Fm* in the *acs1* cultures was generally lower than in the wild-type, with significant decreases at 48, 72, and 144 h (*t*-test, *p* < 0.05) by 4.7%, 8.5%, and 21.6%, respectively (Fig. 4b). These measurements were obtained after dark-grown cells were briefly re-illuminated for *Fv/Fm* analysis, suggesting that the PSII apparatus in *acs1* cells after heterotrophic or mixotrophic growth is more vulnerable to light stress that possibly reflects a high metabolic demand and/or an altered redox balance [28].

Interestingly, the heterotrophically grown *ack2* cultures exhibited significantly higher PSII efficiency than the wild-type upon re-illumination. It maintained significantly higher *Fv/Fm* values at 24, 48, 72, and 96 h (*t*-test, *p* < 0.05) by 6.5%, 6.4%, 6.9%, and 17.6%, respectively (Fig. 4b). The higher *Fv/Fm* values upon re-illumination suggest that loss of *ACK2* does not impair the PSII system during heterotrophic growth.

### 3.5. Real-time respiration profiles of the ACS1 and ACK2 mutants

To better understand energy metabolism in these mutants, we measured real-time oxygen consumption and the respiratory oxygen consumption rate (OCR) of mixotrophic and heterotrophic cultures using the Resipher platform. Under mixotrophic conditions, the *acs1* mutant showed a reduced respiratory activity compared with the wild type. Oxygen depletion proceeded at a slower rate, and the peak OCR was lower than that in the wild-type (Fig. 5a).These data suggest that, under mixotrophic conditions, the *acs1* mutant faces a limitation on maximal respiratory output. This is consistent with previous reports that mitochondrial and chloroplast functions are closely linked in *C. reinhardtii* [12, 20], supporting coordination between respiration, photosynthetic electron transport, and cellular redox state balance. Under heterotrophic conditions, however, the respiratory profile of the *acs1* mutant closely resembled that of the wild-type (Fig. 5c,d), suggesting that loss of *ACS1* does not strongly impair respiration in the dark under the conditions tested.

**Figure 5.**
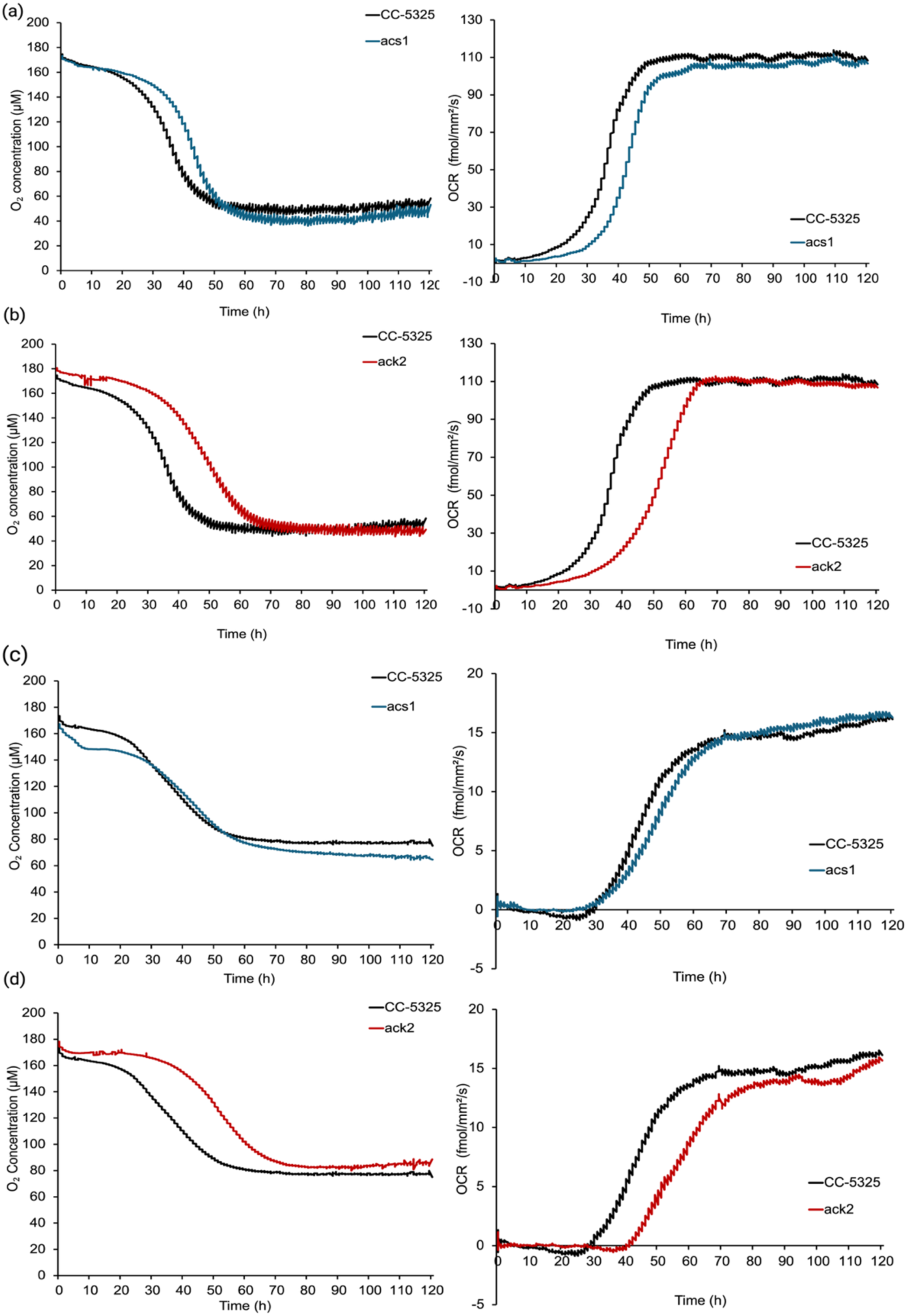
Real-time respiratory profiles of *acs1* and *ack2* mutants under mixotrophic or heterotrophic conditions. Oxygen concentrations (μM; left panels) and oxygen consumption rates (OCR; fmol mm^−2^ s^−1^; right panels) were recorded continuously over 120 h for *acs1* (blue), *ack2* (red), and wild type (CC-5325) (black). Profiles were shown for (a) *acs1* under mixotrophic conditions, (b) *ack2* under mixotrophic conditions, (c) *acs1* under heterotrophic conditions, and (d) *ack2* under heterotrophic conditions. Biological triplicate cultures were maintained in TAP medium at 25 °C in a Resipher 96-well plate to monitor the physiological impact of disruptions in acetate metabolism on respiration.

The *ack2* mutant displayed a more severe respiratory defect. Under mixotrophic conditions, the *ack2* mutant exhibited a noticeable lag in respiratory activity, with peak OCR delayed to *ca.* 65 h compared with *ca.* 50 h in the wild type, along with prolonged retention of dissolved oxygen in the medium (Fig. 5b). Under heterotrophic conditions, this impairment was pronounced, as the *ack2* mutant showed a substantial delay at the onset of rapid oxygen consumption, consistently lower OCR, and higher dissolved oxygen concentrations (Fig. 5d) than wild type across the 120 h time course. At 60 h, the *ack2* mutant showed a 32.4% lower OCR than the wild-type under heterotrophic conditions. Given the mitochondrial localization of ACK2, the respiratory phenotype of the *ack2* knockout is consistent with impaired mitochondrial acetate utilization. By restricting the entry of acetate-derived acetyl-CoA into the TCA cycle, disruption of *ACK2* likely diverts excess carbon into alternative storage sinks.

These respiration results help to explain the observed growth phenotypes. For *acs1*, respiratory defects were observed primarily in the light (Fig. 5a), suggesting that the *acs1* mutation specifically affects how the cell balances energy between acetate catabolism and photosynthesis. In contrast, *ack2* showed delayed and reduced oxygen consumption in both light and dark conditions, suggesting that ACK2 likely catalyzes the primary step for acetate to enter the mitochondrial TCA cycle. Previous metabolic models and flux analyses also suggest that under dark conditions, the ACK/PAT pathway serves as a vital alternative to ACS for maintaining mitochondrial acetyl-CoA [17, 19]. Disrupting this pathway likely limits the supply of acetyl-CoA to the TCA cycle, which could explain the reduced cell density (Fig. 2b). While secondary routes such as the glyoxylate cycle or GABA shunt may allow cells to survive [17], they appear insufficient to fully restore the wild-type respiration or growth rates in the *ack2* mutant.

### 3.6. Lipid and carbohydrate accumulation dynamics

Lipids, particularly TAG, and carbohydrates are primary carbon storage sinks in microalgae. Because acetyl-CoA links acetate catabolism to both energy metabolism and central carbon metabolism, changes in its availability may influence the partitioning of carbon into lipid-and carbohydrate-based storage pools. Given that knockout of the *ACS1* and *ACK2* genes affected photosynthetic and respiratory activities, we measured total lipid, total TAG, and total carbohydrate contents to determine their effect on carbon storage compounds.

Under heterotrophic conditions, total lipid content per cell did not show significant differences between both the *acs1* and *ack2* mutants and the wild-type at either 48 h or 120 h, which correspond to the exponential and stationary phases, respectively (Fig. 6a). A similar pattern was observed under mixotrophic conditions, where total lipid content remained comparable between both mutants and the wild-type strains (Fig. S6a). When evaluated on a biomass dry weight basis, total lipid content also showed no significant differences among the three strains (% DW, Fig. S7a,b).

**Figure 6.**
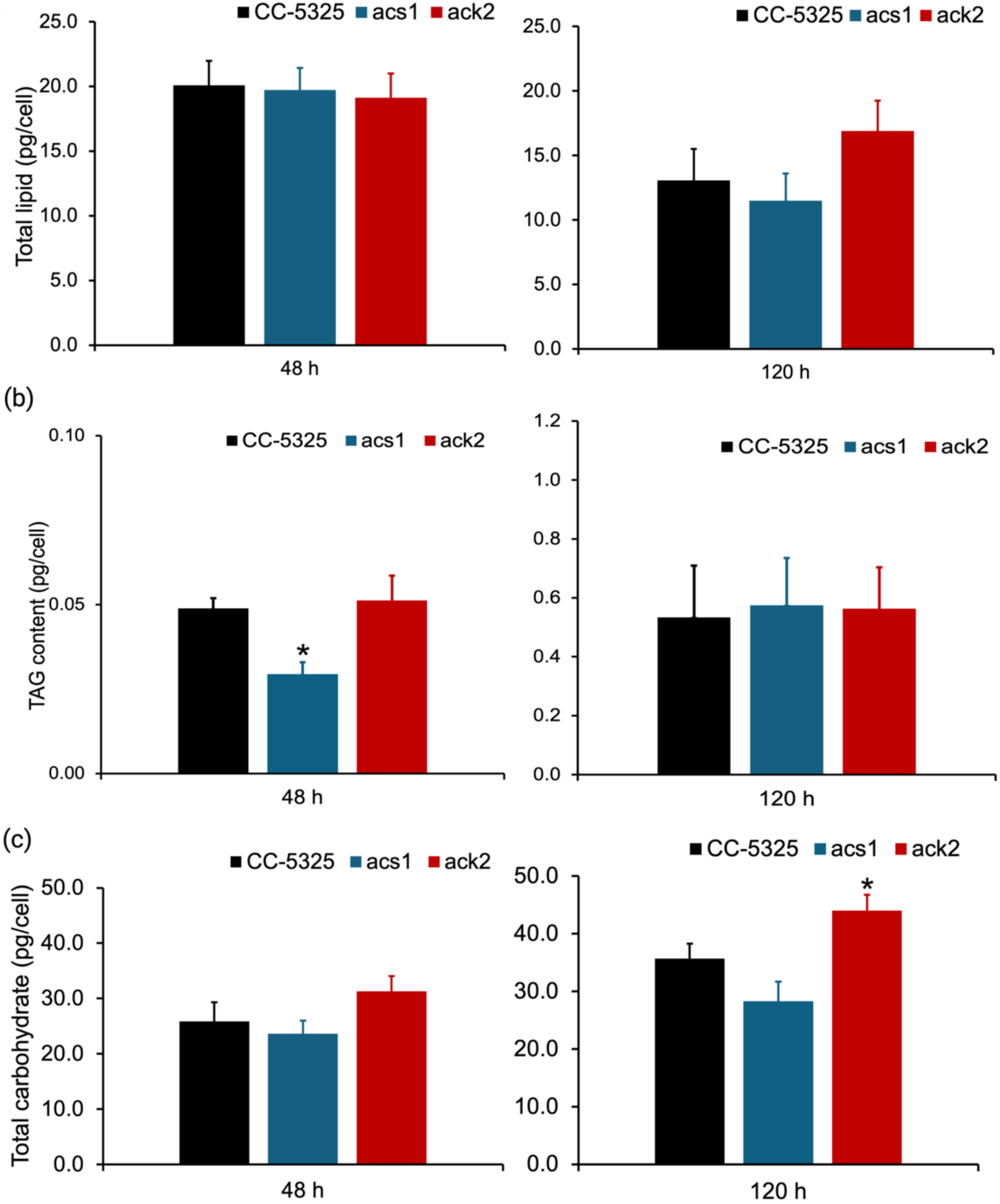
Total lipid, total TAG, and total carbohydrate contents per cell under heterotrophic conditions. Content of (a) total lipid, (b) total TAG, and (c) total carbohydrate in CC-5325 (black), *acs1* (blue), and *ack2* (red) at 48 h and 120 h under heterotrophic conditions. Three biological samples were measured. Error bars represent standard deviations (n=3).Statistical significances between CC-5325 and the mutant strains were determined using a two-tailed Student’s *t*-test, with asterisks (*) indicating a *p*-value < 0.05.

Next, we examined the TAG content, the major storage lipid in cells. Interestingly, *acs1* cells showed a 38.3% lower (*t*-test, *p* < 0.05) TAG content (0.029 pg cell⁻¹) than the wild-type cells (0.047 pg cell⁻¹) under heterotrophic growth at 48 h, whereas *ack2* cells maintained similar TAG levels to the wild-type cells (Fig. 6b). Previously, Fan et al. reported *Chlamydomonas* cells contained 0.1 to 1.3 pg per cell under nutrient-replete mixotrophic conditions containing 20 – 60 mM acetate in the medium, with starch being the dominant storage product [29]. In this study, since we used a lower 17 mM acetate and heterotrophic conditions, the observed values were consistent with low TAG accumulation under nutrient-replete conditions without light. By contrast, TAG levels in *Chlamydomonas* under N-depletion stress during mixotrophic growth were reported to reach *ca.* 40 pg cell⁻¹ [30]. Under mixotrophic conditions, TAG levels remained similar between the wild-type and mutant strains (Fig. S6b), suggesting that disruption of *ACS1* or *ACK2* did not substantially alter TAG storage when both light and acetate were available under nutrient-replete conditions.

TAG is produced via the Kennedy pathway using *de novo*-synthesized acyl-CoA [31, 32] or via membrane lipid remodeling through transesterification of acyl groups [13, 33]. In the *acs1* strain, the loss of cytosolic *ACS1* likely limits the supply of acetate-derived acetyl-CoAs in the early stage of *de novo* TAG biosynthesis, whereas acetyl-CoAs are likely funneled into central carbon metabolism and favor cell division over accumulation of cellular storage compounds (Fig. 1a; 2a). This supports the view that maximizing respiratory flux and cell division often comes at the expense of storage [12, 19]. On the contrary, disruption of the mitochondrial *ACK2* pathway likely reduced mitochondrial acetate utilization and respiratory metabolism, which may lead to limited energy production and slowed cell growth (Fig. 1a; 2a and 5). Under these conditions, assimilated carbon was still directed toward *de novo* fatty acid and TAG biosynthesis, resulting in a wild-type-like TAG content (Fig. 6b).

To determine whether carbon retained in the *ack2* biomass was associated with storage compounds other than lipids, we examined total carbohydrate content. Under mixotrophic conditions, no significant difference was detected between the wild-type and the mutant strains at either the exponential phase or the stationary phase (Fig. S6c). These results show that neither the *acs1* nor the *ack2* mutant showed a change in total lipid accumulation and carbohydrate storage under mixotrophy. Possible explanations are that under mixotrophic conditions, either loss of *ACS1* or *ACK2* has little effect on acetate-derived carbon flux towards carbon storage, or the effects have been buffered by light-driven photosynthesis.

In contrast, a significant difference in total carbohydrate content was detected under heterotrophic growth at 120 h, when the total carbohydrate content in the *ack2* and the wild-type cells reached 44.0 and 35.7 pg cell⁻¹, respectively, representing a 23.3% increase in *ack2* (*t*-test, *p* < 0.05) (Fig. 6c). These results indicate that disruption of *ACK2* promotes an increase in total carbohydrate accumulation under heterotrophic conditions. These data are consistent with the view that loss of mitochondrial *ACK2* reduces respiratory activity and favors carbon accumulation in carbohydrates over TAGs. In corroboration, previous works suggested that in *Chlamydomonas* [34] and another green microalga, *Pseudochlorococcum* sp. [35], carbohydrates such as starch can serve as primary carbon-storage compounds under favorable growth conditions, while TAG accumulation becomes more prominent under stress conditions. The relative partitioning of carbon between starch and TAG accumulation depends on physiological and environmental conditions, such as acetate availability, light, and CO₂, because both storage pathways require carbon skeletons and ATP, while lipid biosynthesis requires more reducing power such as NADPH [36].

## 4. CONCLUSION

This study investigated the roles of *ACS1* and *ACK2* in *C. reinhardtii.* The *acs1* mutant exhibited accelerated growth but reduced early-stage TAG accumulation. In contrast, the *ack2* mutant displayed diminished respiration and a slower growth rate, yet allocated a greater portion of carbon to biomass accumulation, particularly in the form of carbohydrates. These findings demonstrate that *ACS1* and *ACK2* differentially regulate cell growth and carbon storage in *C. reinhardtii*, as proposed in the working model shown in Fig. 7.

**Figure 7.**
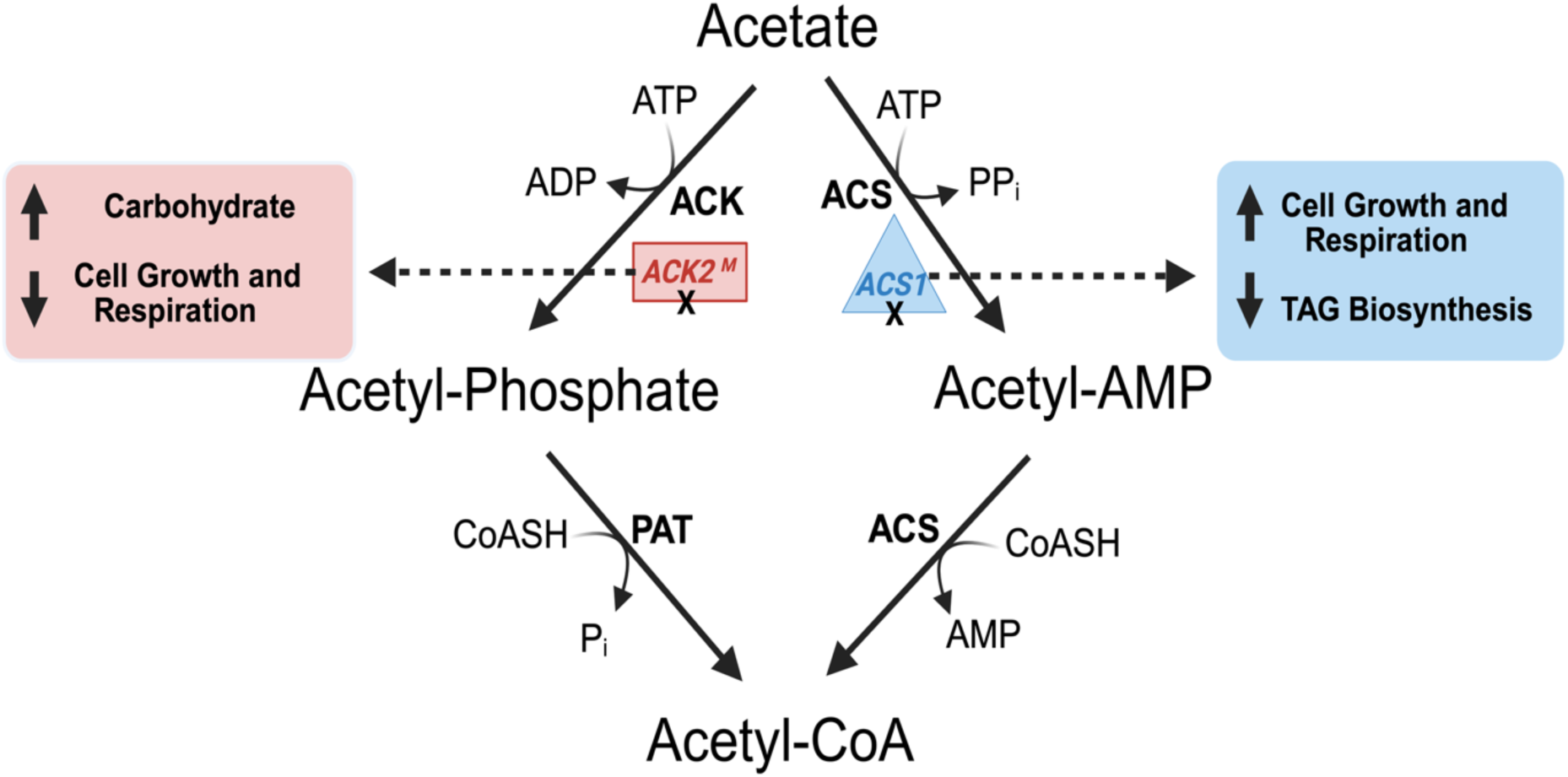
Schematic illustration of acetate catabolism in *C. reinhardtii* under heterotrophic conditions. The diagram outlines the enzymatic routes for converting acetate to the central metabolite acetyl-CoA. Acetate is metabolized via two primary pathways: 1) the ACK-PAT pathway, converting acetate to acetyl-phosphate and subsequently to acetyl-CoA by acetate kinase (ACK), followed by phosphate acetyltransferase (PAT); and 2) the ACS pathway, a two-step reaction converting acetate to acetyl-CoA via an intermediate acetyl-AMP by acetyl-CoA synthetase (ACS). The regulatory impact of gene disruptions is highlighted: loss of cytosolic *ACS1* (blue triangle) results in a metabolic shift favoring increased cell growth and respiration at the expense of TAG biosynthesis, whereas loss of mitochondrial *ACK2* (red rectangle) promotes carbohydrate and biomass storage while reducing growth and respiratory rates. Illustration created with BioRender.com.

This work lays the foundation for integrating reverse genetics with physiological and molecular analyses to investigate acetate metabolism in microalgae. Future studies may incorporate stable-isotope labeling to precisely map carbon fluxes, as well as comparative transcriptomic analyses of the wild type and mutant strains to identify key regulators of carbon metabolism in *Chlamydomonas*. These insights may also be applicable to other microalgal systems and may inform metabolic engineering strategies that couple acetate metabolism with downstream biosynthetic pathways, thereby enhancing the industrial potential of microalgae for the production of biofuel and high-value bioproducts [2, 4].

## ACKNOWLEDGMENTS

A.A. gratefully acknowledges graduate scholarship funding support from King Saud University. This work was partly supported by the Institute of Marine and Environmental Technology (IMET) Angel Investors Program. We thank Dr. Linan Zhang from Qingdao Agricultural University for help with preliminary work associated with the *Chlamydomonas* mutant strains. We thank Dr. Allen Place and Emily Jolly at the Institute of Marine and Environmental Technology for their help with the Resipher experiments, and Yuting Lin for assistance with GC–MS analysis. We also thank Dr. Kyarii Ramarui for help with HPLC measurements.

## AUTHOR CONTRIBUTIONS

**Abdulmajid M. Alrefaie**: Conceptualization (supporting); Data curation (lead); Formal analysis (lead); Investigation (lead); Methodology (lead); Validation (lead); Visualization (lead); Writing – original draft (lead); Writing – review and editing (lead).

**Yi-Ying Lee**: Conceptualization (supporting); Methodology (supporting); Validation (supporting); Investigation (supporting).

**Yantao Li**: Conceptualization (lead); Investigation (supporting); Supervision (lead); Writing – review and editing (lead).

