## Supplementary File for "Characterization of acetate catabolism in *Chlamydomonas reinhardtii* reveals distinct roles for *ACS1* and *ACK2* in regulating cell growth and carbon storage"

### Supplemental materials

Phenotypic screening confirmed the stable integration of the *AphVIII* resistance cassette. When cultured on TAP agar plates supplemented with paromomycin, both mutant strains exhibited robust growth, whereas CC-5325 failed to grow, indicating functional expression of the antibiotic resistance marker (Fig. S1). Additionally, PCR with flanking primers spanning the predicted insertion sites produced larger amplicons in the mutants than in the wild type. The *acs1* mutant yielded a ~4 kb fragment compared to 1175 bp in wild type, and *ack2* also produced a ~4 kb fragment compared to 1536 bp in wild type (Fig. S2). Optimization with GC Buffer and DMSO improved amplification of these GC-rich regions. Lastly, Sanger sequencing confirmed the insertions within internal coding regions: exon 17 of *ACS1* and intron 2 of *ACK2* (Fig. S3).

The combination of antibiotic selection, distinct PCR products, and sequencing of insertional sites provides a consistent picture of loss-of-function phenotypes, consistent with other insertional mutants in *C. reinhardtii* (Li et al. 2016). Overall, the molecular evidence strongly confirms that the target genes are disrupted in the respective mutants.

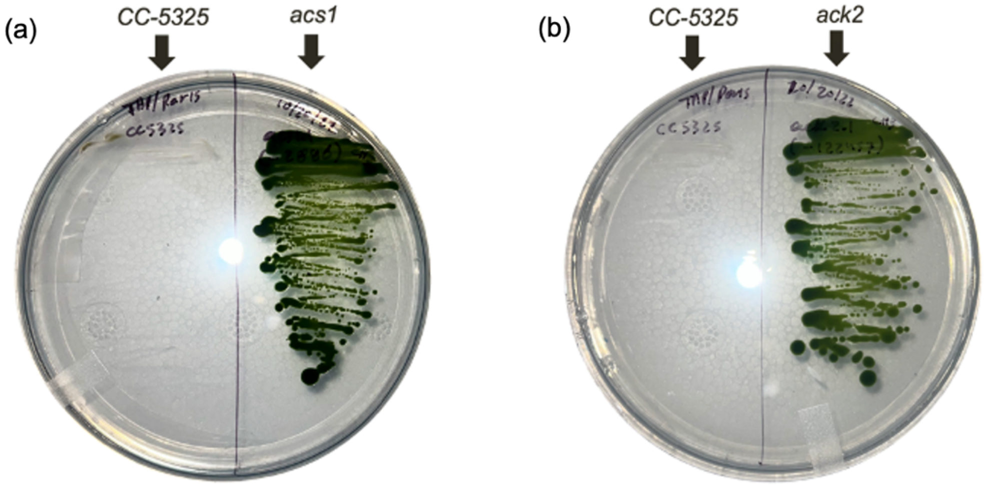


**Supplemental Figure 1. Phenotypic confirmation of *C. reinhardtii* strains of paromomycin resistance. (**a) TAP agar plate supplemented with paromomycin (5 µg $mL^{-1}$) showing the wild type (CC-5325) on the left half and the *acs1* mutant on the right half. (b) TAP agar plate supplemented with paromomycin showing CC-5325 on the left half and the *ack2* mutant on the right half. Robust green colony growth was observed only in the mutants, whereas CC-5325 showed no growth on paromomycin, confirming stable integration and expression of the *AphVIII* resistance cassette in *acs1* and *ack2*.

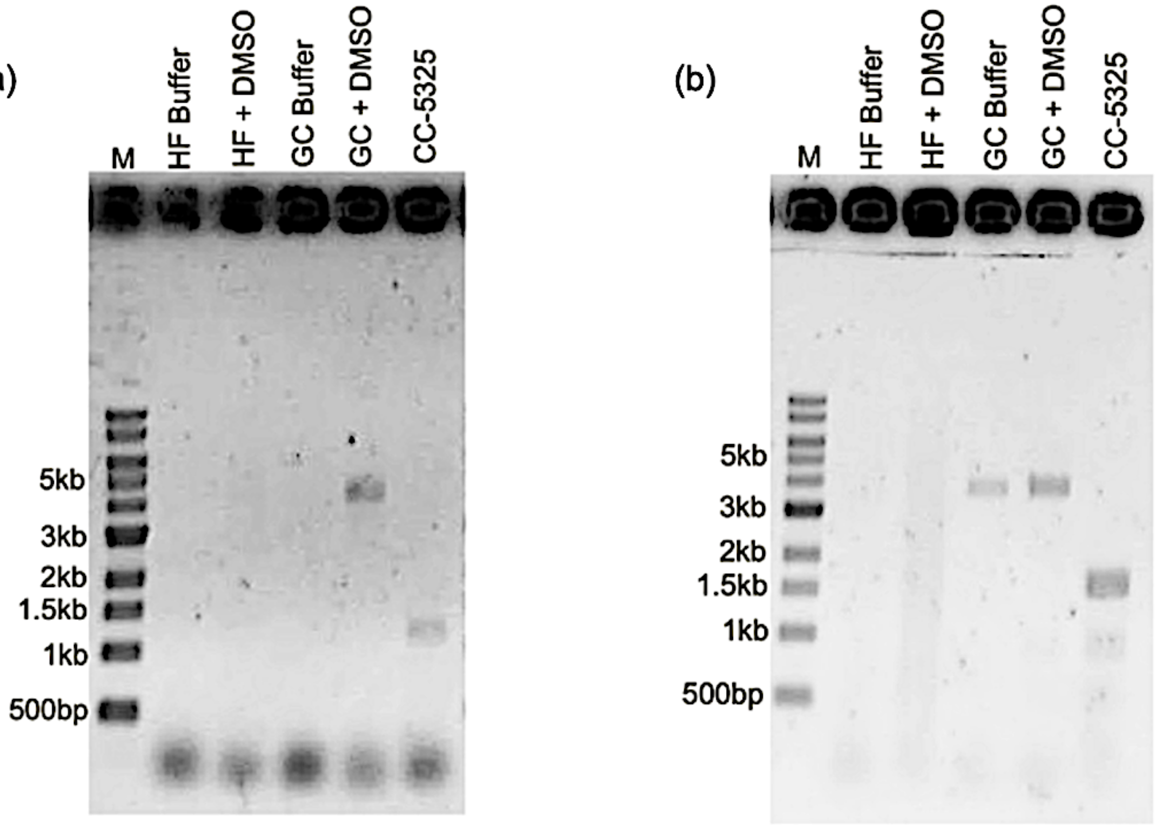


**Supplemental Figure 2. Molecular confirmation of insertional mutations in the *ACS1* and *ACK2* genes.** (a) Agarose gel electrophoresis of PCR products amplified with primers flanking the insertion site in *ACS1* (left). (b) Agarose gel electrophoresis of PCR products amplified with primers flanking the insertion site in *ACK2* (right). For each mutant, reactions were performed under four polymerase buffer conditions (HF buffer, HF buffer + DMSO, GC buffer, GC buffer + DMSO), followed by the control (CC-5325). Lane M, DNA marker with fragment sizes indicated (500 bp–10 kb). In both panels, the mutant lanes showed a ~4-kb band, consistent with insertion of the AphVIII cassette, whereas the CC-5325 lanes displayed shorter fragments (*ACS1*: 1175 bp; *ACK2*: 1536 bp). The distinct band sizes confirmed disruption of the *ACS1* and *ACK2* loci in the *acs1* and *ack2* strains, respectively.

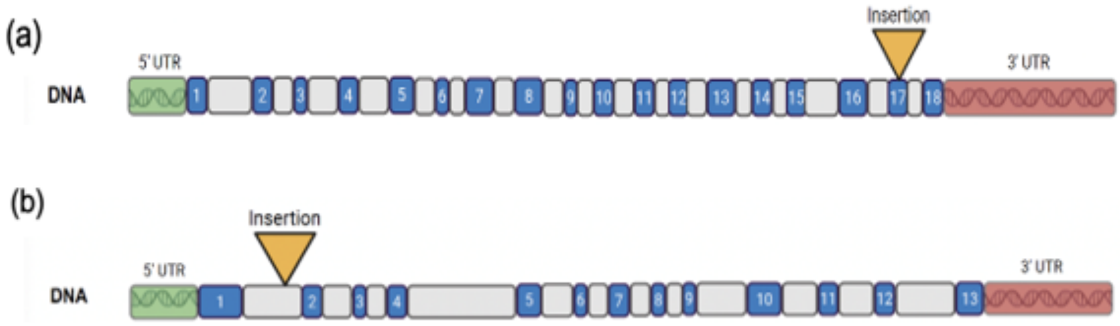


**Supplemental Figure 3. Schematic diagrams of *ACS1* and *ACK2* gene structures and insertion sites.** (a) Map of the *ACS1* locus showing genomic DNA. Blue boxes represented exons, gray boxes represented introns, and green and red segments indicated the 5′ and 3′ untranslated regions (UTRs), respectively. The yellow triangle represented the *AphVIII* cassette insertion within exon 17, indicating disruption of the coding region. (b) Map of the *ACK2* locus showing genomic DNA. Blue boxes represented exons, gray boxes represented introns, and green and red segments indicated the 5′ and 3′ untranslated regions (UTRs), respectively. The yellow triangle marked the *AphVIII* cassette insertion within intron 1. Illustration created with BioRender.com.

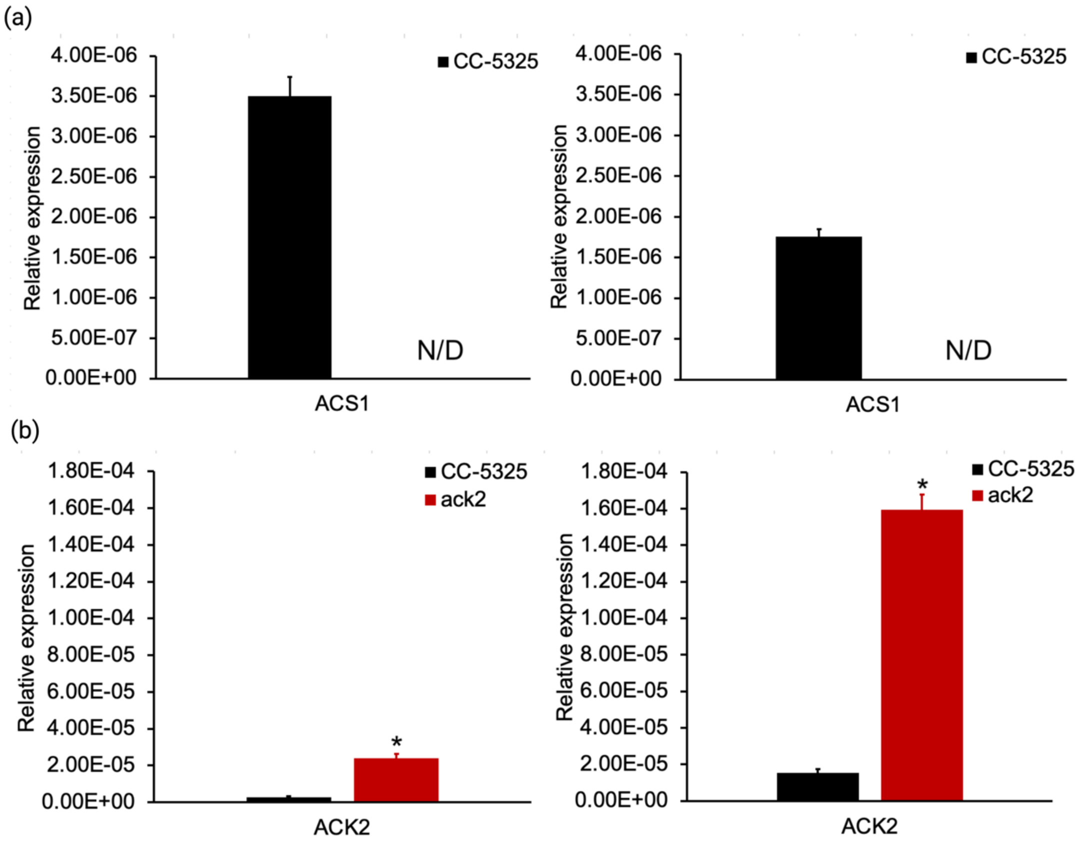


**Supplemental Figure 4. Transcriptional regulation of acetate metabolism in the *acs1* mutant under heterotrophic conditions.** Relative gene expression levels of *ACS1* were determined by RT-qPCR in the CC-5325 (black) and *acs1* mutant strains. Data are shown for the 0 h (left) and 24 h (right) time points. *ACS1* expression was detected in CC-5325 but was non-detectable in the *acs1* mutant at both time points. Expression was normalized to the 18S rRNA gene. Triplicate biological samples were analyzed, and error bars represented standard deviations (SD; n=3). N/D, non-detectable.

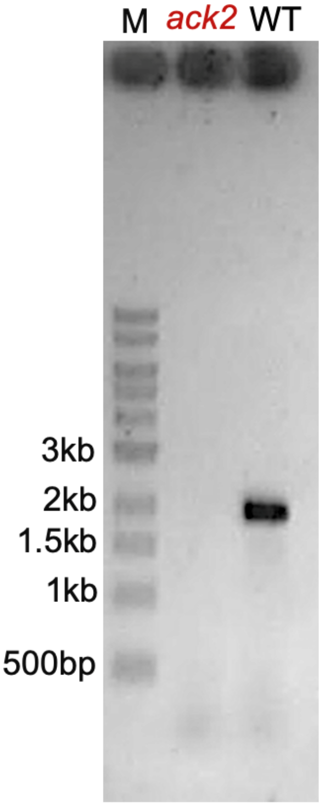


**Supplemental Figure 5. Full-length *ACK2* cDNA amplification in the wild type and the *ack2* mutant.** Agarose gel electrophoresis of RT-PCR products amplified with primers spanning the near full-length *ACK2* transcript, with an expected product size of 1919 bp. A band of the expected size was amplified from CC-5325 but not from the *ack2* mutant. Lane M, DNA marker with fragment sizes indicated (500 bp–3 kb). These results indicate that the insertion in *ack2* disrupts transcript continuity and prevents production of an intact full-length *ACK2* mRNA.

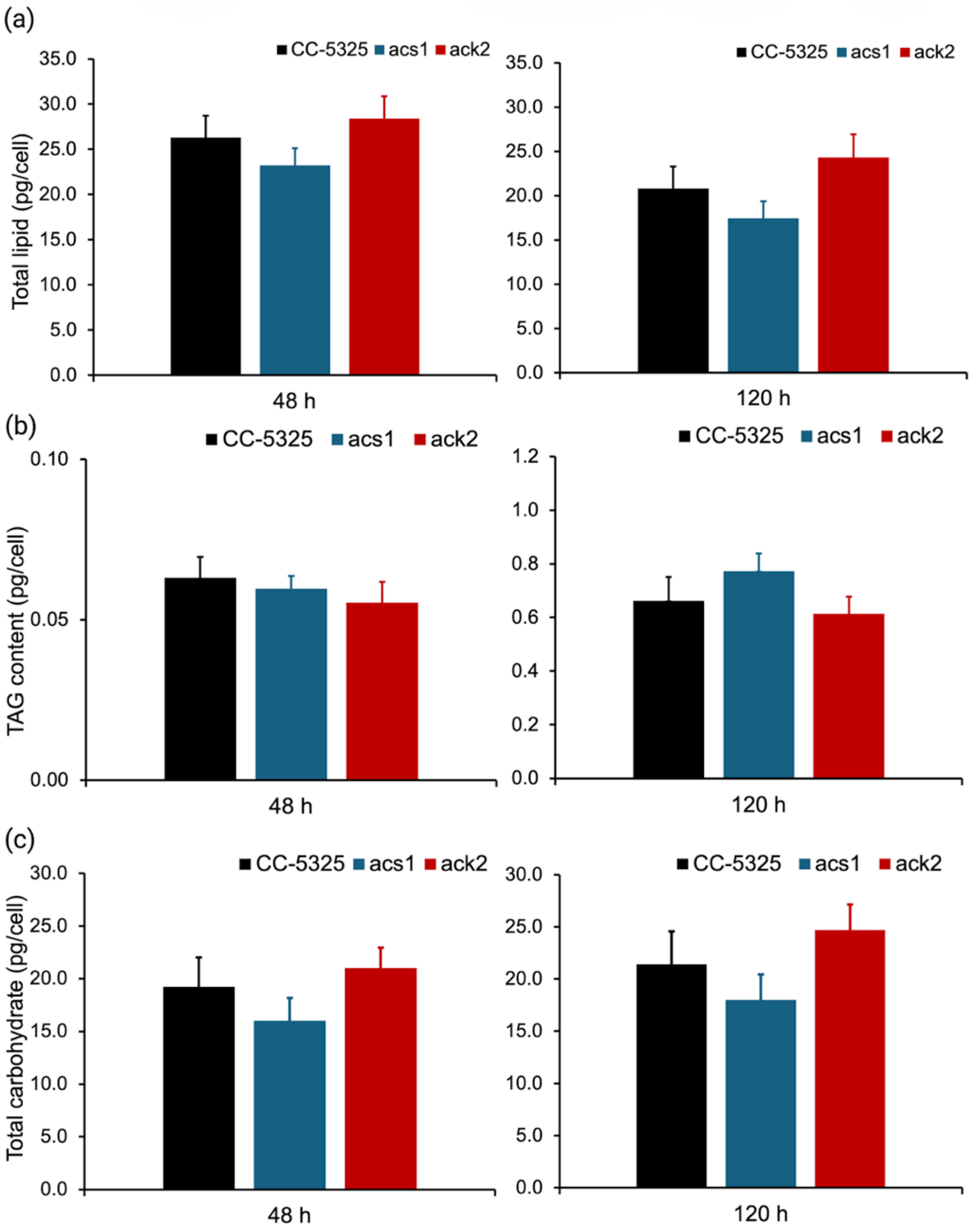


**Supplemental Figure 6. Total lipid, total TAG, and total carbohydrate contents per cell under mixotrophic conditions.** Content of (a) Total lipid, (b) total TAG, and (c) total carbohydrate in CC-5325 (black), *acs1* (blue), and *ack2* (red) at 48 h and 120 h under mixotrophic conditions. Three biological samples were measured. Error bars represent standard deviations (n = 3). Statistical significances between CC-5325 and the mutant strains were determined using a two-tailed Student's *t*-test, with asterisks (*) indicating a $p$-value < 0.05.

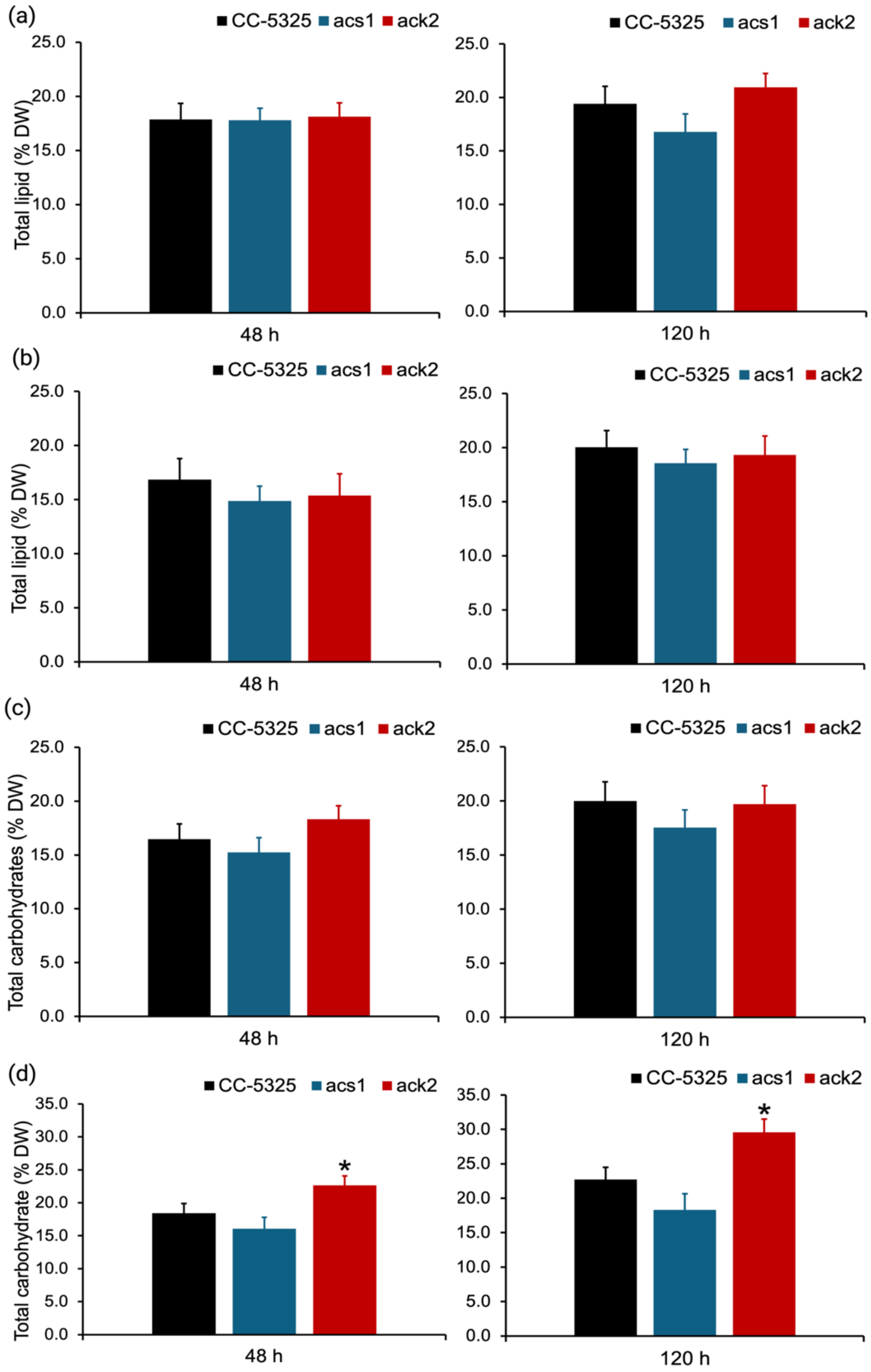


**Supplemental Figure 7. Total lipid and total carbohydrate contents on a biomass dry weight basis under mixotrophic and heterotrophic conditions.** Total lipid content (% dry weight, DW) at 48 and 120 h under (a) mixotrophic and (b) heterotrophic conditions. Total carbohydrate content (% DW) at 48 h and 120 h under (c) mixotrophic and (d) heterotrophic conditions. Three biological samples were measured. Error bars represented standard deviations (n=3). Statistical significances between CC-5325 and *ack2* were calculated using a two-tailed Student's *t*-test, with asterisks (*) indicating a *p*-value < 0.05.

| Protein | Experimentally confirmed[a] | PredAlgo[b] | DeepLoc 2.0[c] |
|---|---|---|---|
| ACS1 | - | Other | Cytosol |
| ACS2 | - | Chloroplast | Chloroplast |
| ACS3 | - | Mitochondrion | Mitochondrion / Cytosol |
| ACK1 | Chloroplast | Chloroplast | Chloroplast / Cytosol |
| ACK2 | Mitochondrion | Mitochondrion | Mitochondrion / Cytosol |
| PAT1 | Mitochondrion | Mitochondrion | Mitochondrion |
| PAT2 | Chloroplast | Chloroplast | Chloroplast |

[a]Localization based on experiments
[b]Localization based on *in silico* predictions from PredAlgo
[c]Localization based on *in silico* predictions from DeepLoc 2.0

**Supplemental Table 1. Prediction of subcellular localization for proteins involved in acetate metabolism.** The table summarized experimentally supported (a) localizations and *in silico* predictions from PredAlgo (b) and DeepLoc 2.0 (c) for acetyl-CoA synthetases ACS1–3, acetate kinases ACK1–2, and phosphate acetyltransferases PAT1–2. Literature-based localizations were compiled from published proteomic and biochemical studies. PredAlgo and DeepLoc 2.0 outputs indicated whether each protein was targeted to the cytosol, chloroplast, mitochondria, or multiple compartments.
